# The dynamic interplay of host and viral enzymes in type III CRISPR-mediated cyclic nucleotide signalling

**DOI:** 10.1101/2020.02.12.946046

**Authors:** Januka S. Athukoralage, Shirley Graham, Christophe Rouillon, Sabine Grüschow, Clarissa M. Czekster, Malcolm F. White

## Abstract

Cyclic nucleotide second messengers are increasingly implicated in prokaryotic anti-viral defence systems. Type III CRISPR systems synthesise cyclic oligoadenylate (cOA) upon detecting foreign RNA, activating ancillary nucleases that can be toxic to cells, necessitating mechanisms to remove cOA in systems that operate via immunity rather than abortive infection. Previously, we demonstrated that the *Sulfolobus solfataricus* type III-D CRISPR complex generates cyclic tetra-adenylate (cA_4_), activating the ribonuclease Csx1, and showed that subsequent RNA cleavage and dissociation acts as an “off-switch” for the cyclase activity (Rouillon *et al.*, 2018). Subsequently, we identified the cellular ring nuclease Crn1, which slowly degrades cA_4_ to reset the system, and demonstrated that viruses can subvert type III CRISPR immunity by means of a potent anti-CRISPR ring nuclease variant. Here, we present a comprehensive analysis of the dynamic interplay between these enzymes, governing cyclic nucleotide levels and infection outcomes in virus-host conflict.

## Introduction

CRISPR systems are widespread in archaea and bacteria, providing adaptive immunity against invading mobile genetic elements (MGE) (Sorek *et al.*, 2013, Makarova *et al.*, 2020). Type III CRISPR systems (Figure 1) are multi-functional effector proteins that have specialised in the detection of foreign RNA (Tamulaitis *et al.*, 2017, Zhu *et al.*, 2018). The large subunit, Cas10, harbours two enzyme active sites that are activated by target RNA binding: a DNA-cleaving HD nuclease domain (Samai *et al.*, 2015, Elmore *et al.*, 2016, Estrella *et al.*, 2016, Kazlauskiene *et al.*, 2016) and a cyclase domain for cyclic oligoadenylate (cOA) synthesis (Kazlauskiene *et al.*, 2017, Niewoehner *et al.*, 2017, Rouillon *et al.*, 2018). The third enzymatic activity of type III systems is situated in the Cas7 subunit of the complex, which cleaves bound RNA targets and in turn regulates Cas10 enzymatic activities (Tamulaitis *et al.*, 2014, Rouillon *et al.*, 2018, Johnson *et al.*, 2019, Nasef *et al.*, 2019). The cyclase domain polymerises ATP into cOA species consisting of between 3-6 AMP subunits (denoted cA_3_, cA_4_ etc.), in varying proportions (Kazlauskiene *et al.*, 2017, Niewoehner *et al.*, 2017, Rouillon *et al.*, 2018, Grüschow *et al.*, 2019, Nasef *et al.*, 2019). cOA second messengers activate CRISPR ancillary nucleases of the Csx1/Csm6, Can1 (CRISPR associated nuclease 1) and NucC families, which drive the immune response against MGEs (Kazlauskiene *et al.*, 2017, Niewoehner *et al.*, 2017, Rouillon *et al.*, 2018, Grüschow *et al.*, 2019, McMahon *et al.*, 2019, Lau *et al.*, 2020). To date, cA_4_ appears to be the most widely used signalling molecule by type III CRISPR systems (Grüschow *et al.*, 2019). The ribonuclease activity of Csx1/Csm6 is crucial for the clearance of MGEs (Hatoum-Aslan *et al.*, 2014, Foster *et al.*, 2019, Grüschow *et al.*, 2019), particularly when viral genes are transcribed late in infection, at low levels or mutated (Hatoum-Aslan *et al.*, 2014, Jiang *et al.*, 2016, Rostol & Marraffini, 2019).

**Figure 1.**
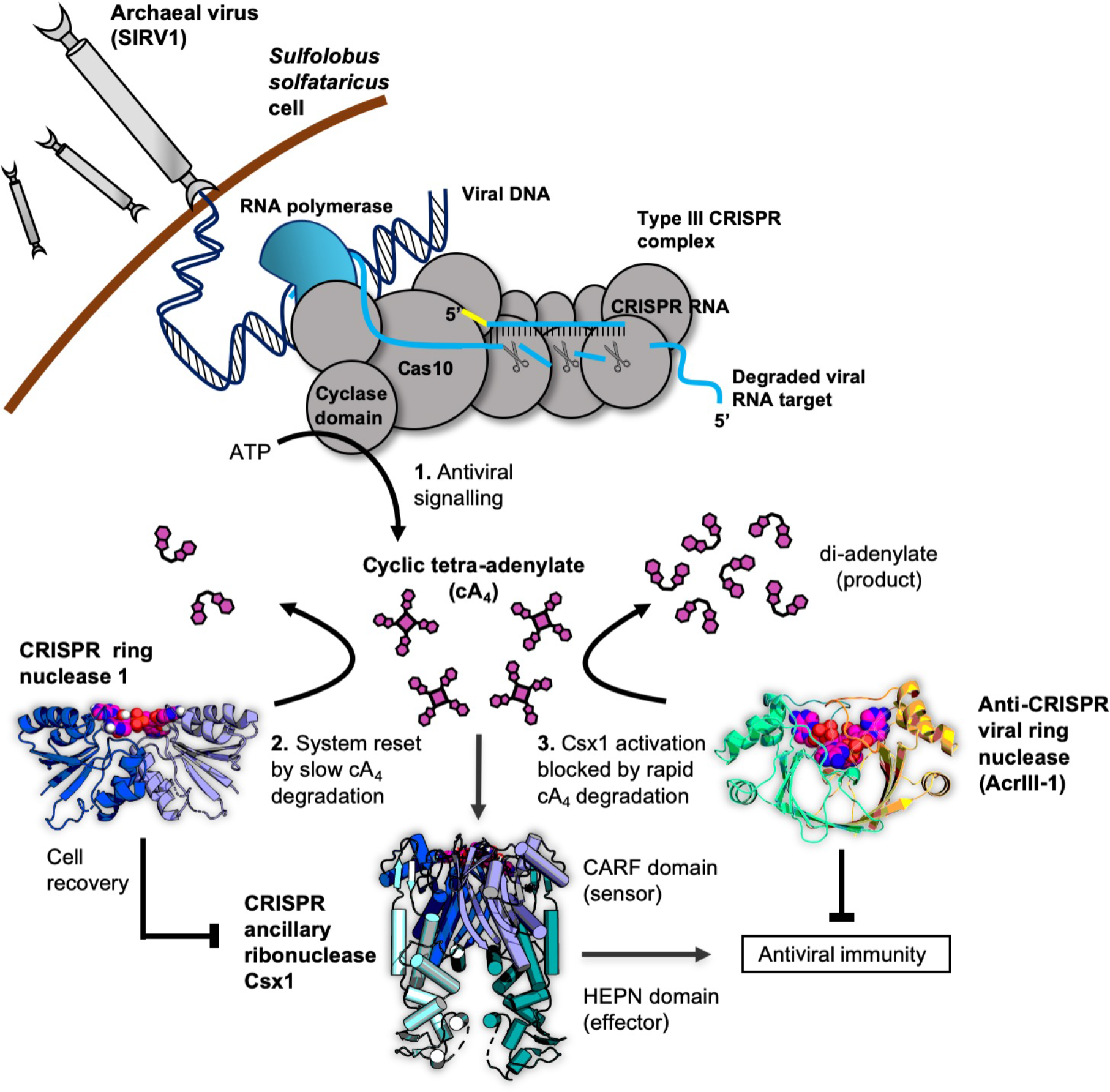
Cartoon of type III CRISPR cyclic nucleotide signalling and defence in *Sulfolobus solfataricus*. The Cas10 subunit of the type III CRISPR complex synthesises cyclic tetra-adenylate (cA_4_) when viral RNA transcripts are detected. Target RNA cleavage shuts-off cA_4_ synthesis. cA_4_ binds to CARF (CRISPR associated Rosmann Fold) domain of CRISPR ancillary nuclease Csx1 and allosterically activates its HEPN (Higher Eukaryotes and Prokaryotes Nucleotide binding) domain, which degrades RNA non-specifically within the cell. Extant cA_4_ is degraded slowly by CRISPR ring nucleases (Crn1 family) which likely facilitate cell recovery after clearing the virus. Viral anti-CRISPR ring nucleases (AcrIII-1 family) degrade cA_4_ rapidly to stop activation of ancillary defence enzymes such as Csx1 and supress antiviral immunity.

In our previous study, we demonstrated that the type III-D system from *Sulfolobus solfataricus* synthesises predominantly cA_4_, which activates the CRISPR ancillary ribonuclease Csx1. We examined the first regulatory step in cOA synthesis in detail and demonstrated that target RNA cleavage and dissociation from the complex shut-off cOA synthesis (Rouillon *et al.*, 2018). Since CRISPR ancillary nucleases degrade nucleic acids non-specifically, cellular as well as viral targets are destroyed. Collateral cleavage of self-transcripts by a Csm6 enzyme has previously been shown to result in cell growth arrest (Rostol & Marraffini, 2019). Therefore, in addition to regulating the synthesis of cOA, cells need a mechanism to remove extant cOA if they are to return to normal growth. To solve this problem, *S. solfataricus* encodes CRISPR associated ring nuclease 1 (Crn1) family enzymes (Athukoralage *et al.*, 2018). Crn1 enzymes slowly degrade cA_4_ to yield di-adenylate products incapable of activating Csx1. In other species Csm6 proteins have evolved catalytic CARF domains capable of degrading cA_4_, thereby acting as their own “off-switches” to their RNase activity (Athukoralage *et al.*, 2019, Jia *et al.*, 2019). Unsurprisingly, archaeal viruses and bacteriophage have co-opted this regulatory strategy in order to subvert type III CRISPR defence. Many archaeal viruses and bacteriophage encode a ring nuclease anti-CRISPR (AcrIII-1), unrelated to Crn1, which neutralises the type III response by rapidly degrading cA_4_ to prevent ancillary nuclease activation (Athukoralage *et al.*, 2020).

It is clear that the cA_4_ antiviral second messenger is at the centre of a network of interactions that are crucial for effective type III CRISPR defence against MGE. Here, we show that detection of even a single molecule of invading RNA leads to a large signal amplification by flooding the cell with cA_4_ that in turn activates the non-specific degradative ribonuclease Csx1. We explore how a cellular ring nuclease can return the cell to a basal state and how viruses can subvert the system. By quantifying and modelling the equilibria and reactions that take place in the arena of type III CRISPR defence, we build a comprehensive model of this dynamic, life or death process.

## RESULTS

Whilst the control of cOA synthesis by target RNA binding and cleavage is now understood reasonably well, the full implications of cOA generation in a virally-infected cell are not. This requires a detailed knowledge of the levels of cOA produced, consequences for antiviral defence enzymes and the effects of cOA degrading enzymes from cellular and viral sources. These were the aims of our study.

### Signal amplification on cA_4_ production

We first investigated the extent of signal amplification that occurs in a cell from detection of a single viral RNA and generation of the cA_4_ second messenger. Using the *S. solfataricus* type III-D CRISPR effector, we varied the concentration of target RNA and quantified the resultant cA_4_ production. As previously observed (Rouillon *et al.*, 2018), increasing the target RNA concentration resulted in increased cA_4_ production (Figure 2). Quantification of the concentration of cA_4_ generated was accomplished by using α-^32^P-ATP and quantification of products using a phosphorimager in comparison to standards (Figure 2-figure supplement 1), as described in the methods. We observed that approximately 1000 molecules (980 ± 24) of cA_4_ were generated per molecule of RNA, over a range of 10-100 nM target RNA (Figure 2). When a poorly-cleavable target RNA species containing phosphorothioates was used as the substrate, the amount of cA_4_ generated increased approximately 3-fold (3100 ± 750, Figure 2), confirming the important role of RNA cleavage for deactivation of the cyclase domain (Rouillon *et al.*, 2018, Nasef *et al.*, 2019).

**Figure 2.**
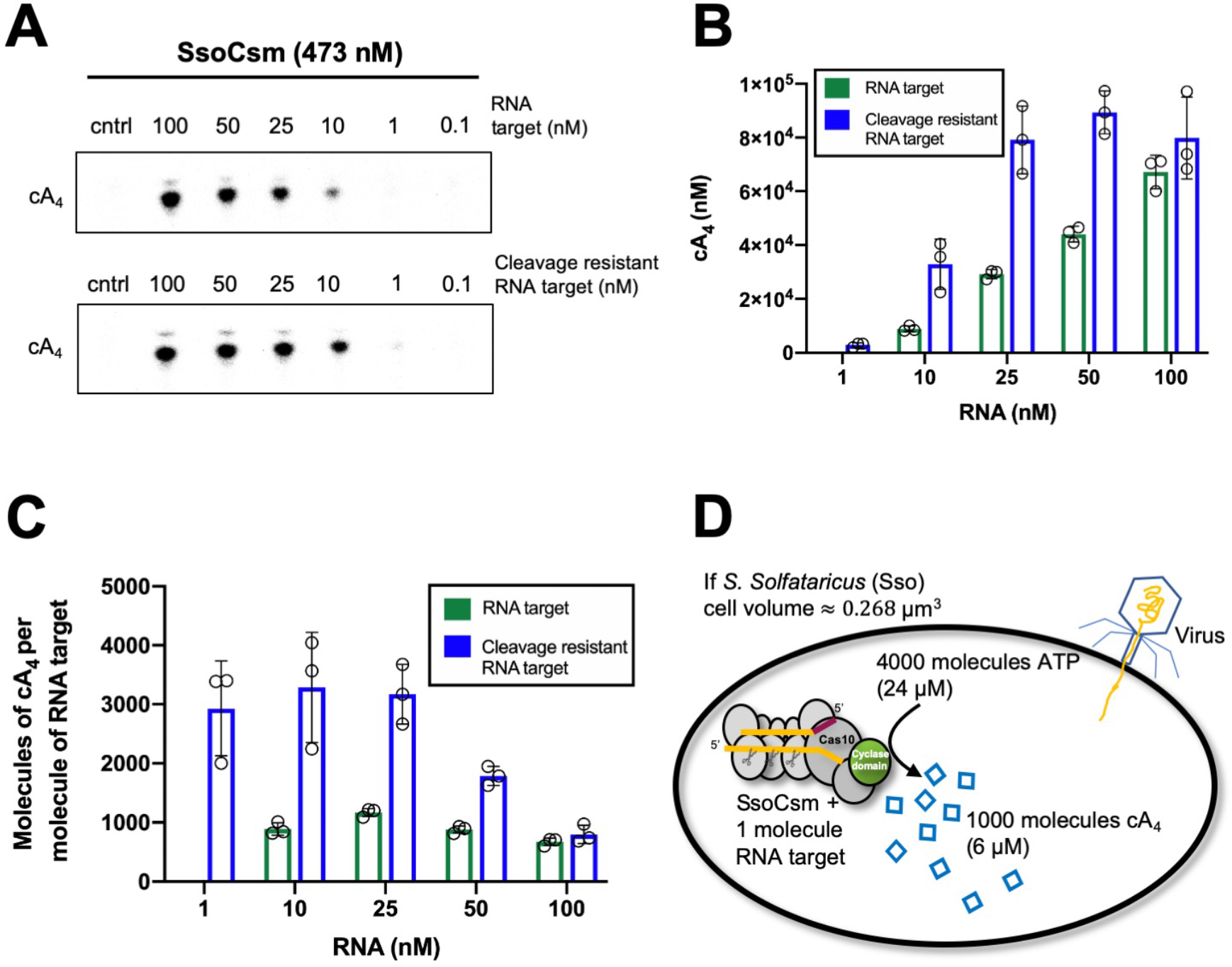
Approximately 1000 molecules cA_4_ are made per molecule of RNA target. **A.** Upper panel shows phosphorimages of thin-layer chromatography of cyclic tetra-adenylate (cA_4_) made by *S. solfataricus* (Sso) Csm complex (470 nM carrying the CRISPR RNA A26) across a range of RNA target concentrations (0.1, 1, 10, 25,100 nM) complementary to the A26 CRISPR RNA at 70 °C. Lower panel shows cA_4_ synthesised with a cleavage resistant (phosphorothioate) form of the RNA target. **B**. Bar graph of the concentration of cA_4_ generated with increasing cleavable and cleavage-resistant RNA target generated by quantifying the densiometric signals from A, with an α-^32^P-ATP standard curve (Figure 2-figure supplement 1). Error bars indicate the standard deviation of the mean of three technical replicates, with individual data points shown as clear circles. No data is shown for 1 nM cleavable RNA target as cA_4_ generated was below detection limits. **C**. Bar chart quantifying the number of molecules of cA_4_ generated per molecule of cleavable or cleavage resistant target RNA across a range of RNA target concentrations. On average SsoCsm synthesised 980 ± 24 and 3100 ± 750 molecules of cA_4_ per molecule of cleavable and cleavage resistant target RNA, respectively. **C**. Cartoon depicting the cellular implications of ~1000 molecules of cA_4_ generated per molecule of RNA target, which in *S. solfataricus* would equate to ~6 μM cA_4_ within the cell.

Given that *S. solfataricus* cells are cocci with a diameter of approximately 0.7 μm, the volume of an average cell can be calculated as approximately 0.3 fL (by comparison, *E. coli* has a cell volume of 1 fL (Kubitschek & Friske, 1986)). Using Avogadro’s number, 1000 molecules equates to an intracellular concentration of 6 μM cA_4_ in *S. solfataricus*. Thus, detection of one viral RNA in the cell would result in the synthesis of 6 μM cA_4_, 10 RNAs – 60 μM, etc. The upper limits of cA_4_ generation could be defined by the number of viral target RNAs present, the number of type III effectors carrying a crRNA matching that target, or even conceivably the amount of ATP available for cA_4_ generation.

### Kinetic parameters of the Csx1 ribonuclease

The cA_4_ second messenger binds to CARF family proteins to elicit an immune response. To understand the concentration of cA_4_ required to activate an antiviral response, we determined the dissociation constant of the major ancillary ribonuclease Csx1 for the cA_4_ activator. Using radioactive cA_4_, we titrated an increasing concentration of Csx1 protein and subjected the mixture to native gel electrophoresis (Figure 3A, B). cA_4_ was bound by Csx1 with a dissociation constant of 430 ± 40 nM. Thus, even one viral target RNA detected by the type III CRISPR system should generate enough cA_4_ (6 μM) to fully activate the Csx1 ribonuclease for defence. We proceeded to estimate the binding affinity of a ribonuclease-deficient Csx1 variant for its RNA target, yielding an apparent dissociation constant of approximately 5 μM (Figure 3C), and determined the single-turnover kinetic constant for cA_4_-activated RNA cleavage by Csx1 as 5.8 ± 0.6 min^−1^ (Figure 4).

**Figure 3:**
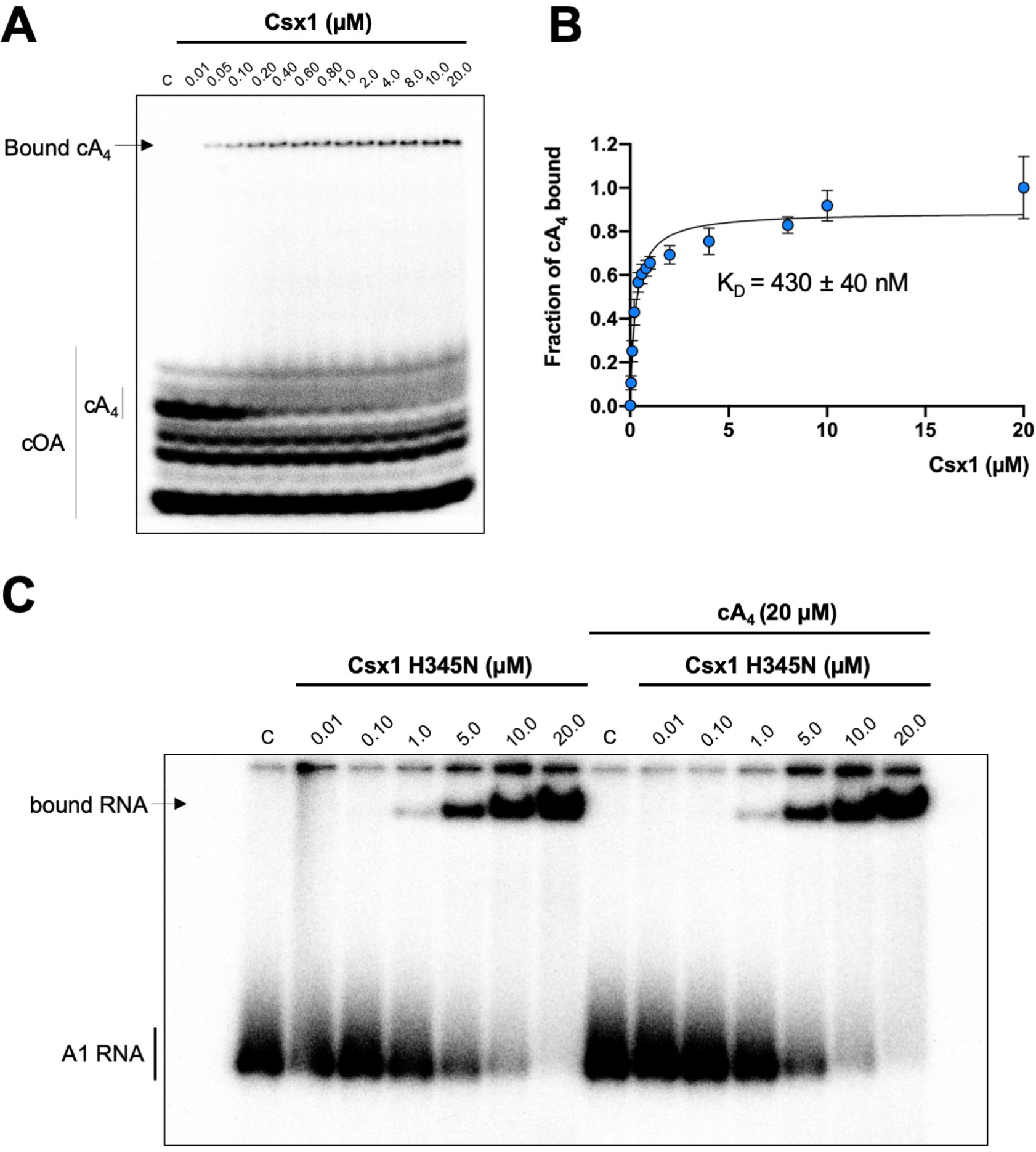
Csx1 binds cA_4_ with high affinity and RNA with relatively low affinity. **A.** Phosphorimage of native gel electrophoresis visualising cA_4_ (20 nM) binding by Csx1 (concentrations as indicated in the figure). **B**. Plot of fraction of cA_4_ bound by Csx1. Error bars indicate the standard deviation of the mean of four technical replicates and the data is fitted to the equation (Fraction cA_4_ bound = Bound_max_ / (1+ (K_D_/ [SsoCsx1])); Bound_max_ =1). **C**. Phosphorimage of native gel electrophoresis visualising A1 substrate RNA binding by Csx1 H345N protein dimer in the absence (left hand-side) or presence (right hand-side) of unlabelled cA_4_ (20 μM). The image shown is representative of three technical replicates. Control c – RNA alone.

**Figure 4.**
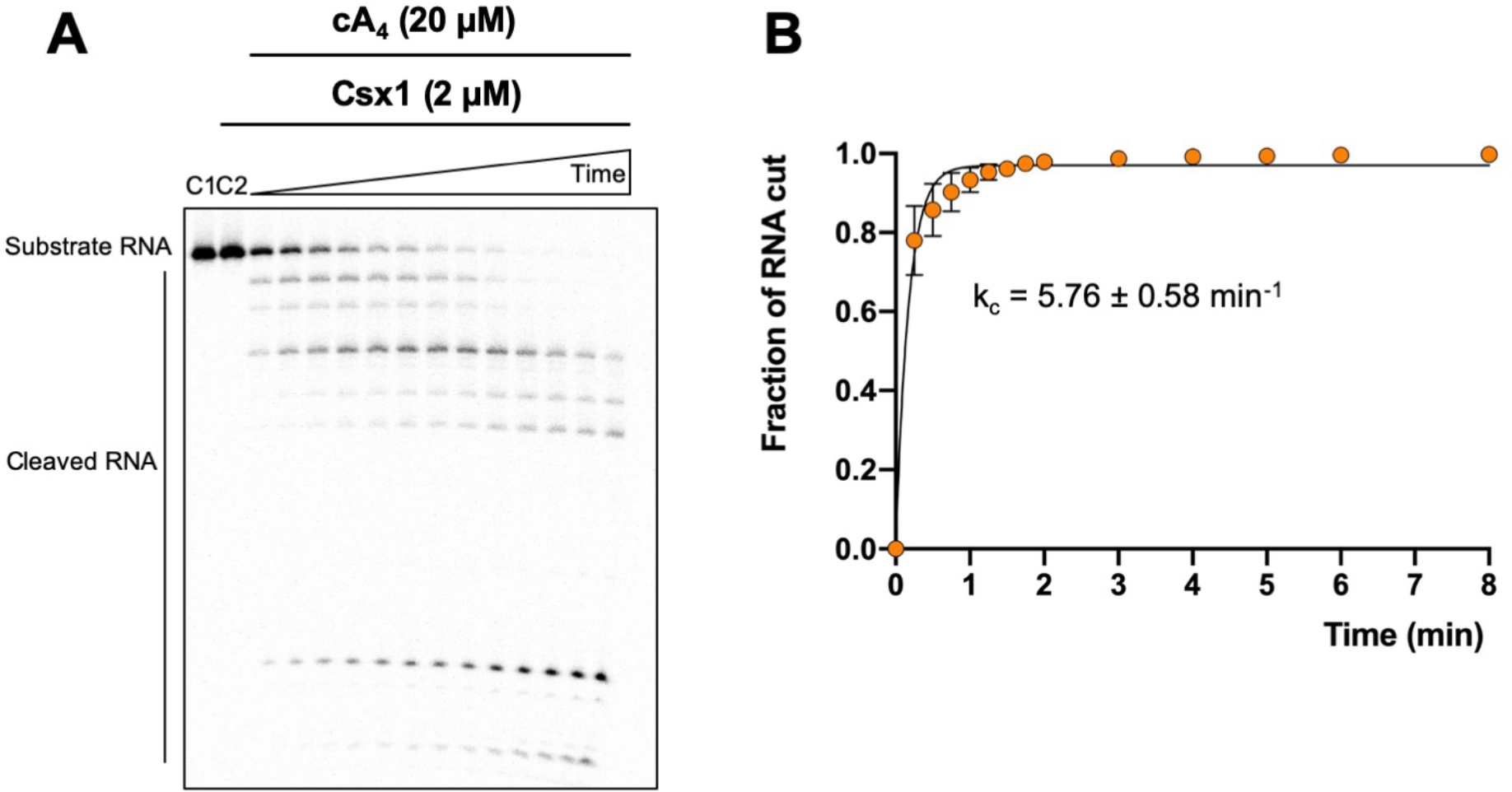
Degradation of RNA by Csx1. **A.** Phosphorimage of denaturing gel electrophoresis visualising A1 RNA (50 nM) cleavage by an excess of Csx1 (2 μM dimer) at 50 °C. Controls: C1 – RNA alone; C2 – reaction in the absence of cA_4_ for 8 min. **B**. Plot of fraction of RNA cut by Csx1 over time. The data is fitted to an exponential equation and error bars show the standard deviation of the mean of three technical replicates.

### Kinetic and equilibrium constants of the ring nucleases Crn1 and AcrIII-1

We have previously established that Crn1 cleaved cA_4_ at a rate of 0.089 ± 0.003 min^−1^ at 50 °C, while AcrIII-1 cleaved cA_4_ at a rate of 5.4 ± 0.38 min^−1^, about 60-fold faster (Athukoralage *et al.*, 2020). The difference in reaction rates probably reflects the different roles of the two enzymes, with Crn1 working in conjunction with the type III CRISPR defence and AcrIII-1 opposing it. To quantify the interaction between ring nucleases and cA_4_, we titrated radioactively labelled cA_4_ with either Crn1 or AcrIII-1 and visualised cA_4_ binding by phosphorimaging following native gel electrophoresis. Crn1 bound cA_4_ with an apparent dissociation constant of ~50 nM, while the inactive H47A variant of AcrIII-1 bound cA_4_ with an apparent dissociation constant of ~25 nM (Figure 5). Thus, both ring nucleases bound cA_4_ about 10-fold more tightly than Csx1.

**Figure 5.**
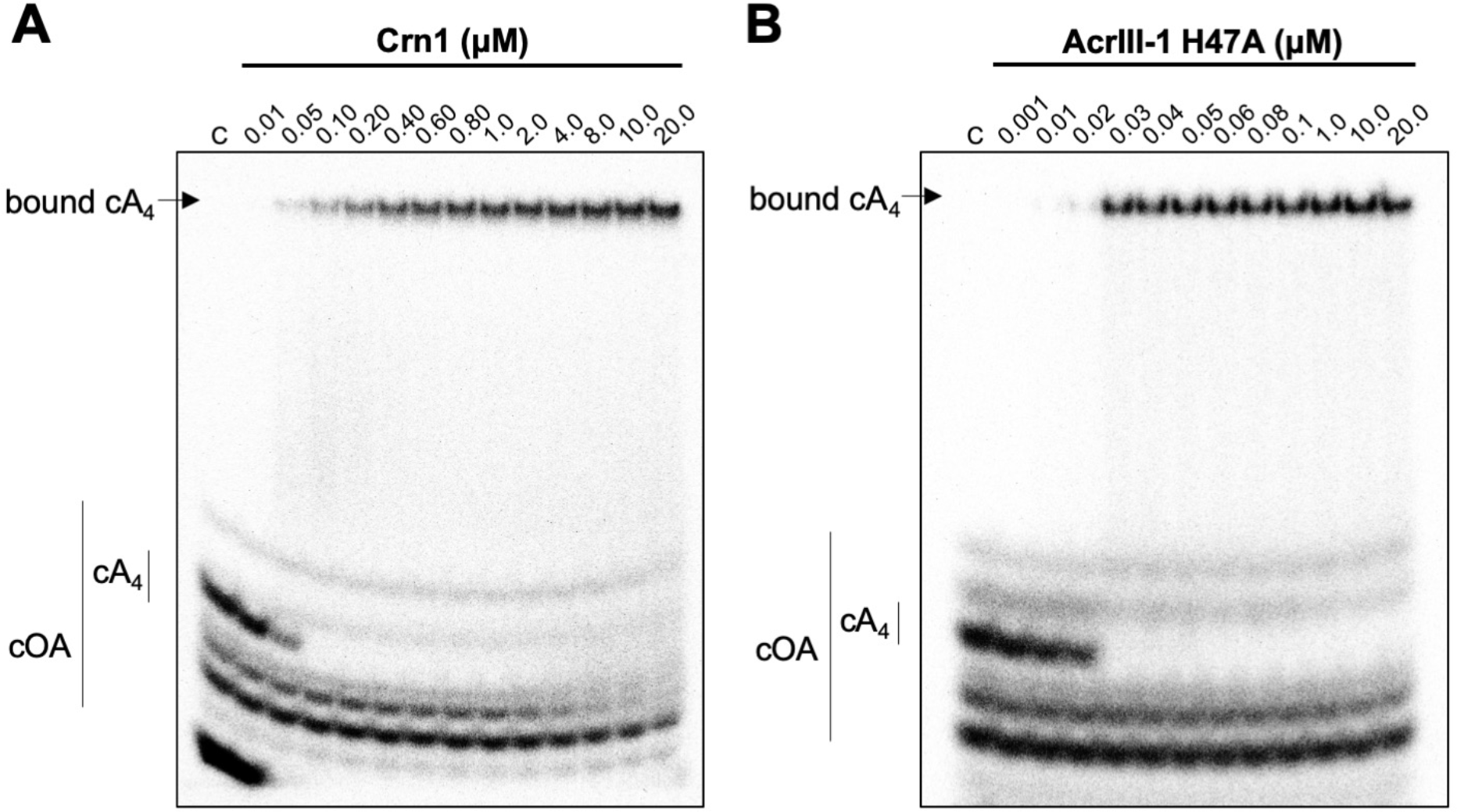
Crn1 and AcrIII-1 bind cA_4_ with high affinity. Phosphorimages of native gel electrophoresis visualising radiolabelled cyclic oligoadenylate (cOA) binding by (**A**) Crn1 (**B**) and catalytically inactive AcrIII-1 (SIRV1 gp29 H47A). Crn1 binds cA_4_ (10 nM) with an apparent dissociation constant of approximately 50 nM, whereas AcrIII-1 binds cA_4_ with an apparent dissociation constant of approximately 25 nM. The images shown are representative of three technical replicates. Control c – cA_4_ alone.

### Kinetic modelling of the antiviral signalling pathway and its regulation by cA_4_ degrading enzymes

We entered the experimentally determined kinetic and equilibria parameters into the KinTek Global Kinetic Explorer software package and generated a model to simulate RNA degradation by Csx1 and the effects of ring nucleases over time (Figure 6A and Table 1). We first examined RNA cleavage by Csx1 in the presence of 6, 60 or 600 μM cA_4_ (equivalent to low, medium and high levels of infection). In all cases, the input RNA (100 μM) was almost fully cleaved by 48 h, suggesting that unregulated Csx1 activity could result in cellular stress (Figure 6-figure supplement 1). Under these conditions, Csx1 was fully activated regardless of the simulated level of infection due to its high affinity for cA_4_ – a situation that might not be favourable *in vivo*. Next we evaluated the effect of the cellular ring nuclease Crn1 in the model. In agreement with biochemical assays in which Crn1 was able to deactivate Csx1 by degrading low levels of cA_4_ (Athukoralage *et al.*, 2020), in our simulations 1 μM Crn1 effectively degraded 6 μM cA_4_ corresponding to a single RNA target to deactivate Csx1 (Figure 6B). In contrast, when challenged with 60 μM cA_4_, Crn1 deactivated Csx1 more slowly. At the highest concentration of cA_4_ (600 μM), Crn1 could not degrade the activator in time to prevent Csx1 cleaving all the RNA (Figure 6C). Thus, addition of a ring nuclease activity allows the cell to respond to different levels of infection, and therefore cA_4_ concentration, in different ways.

**Figure 6.**
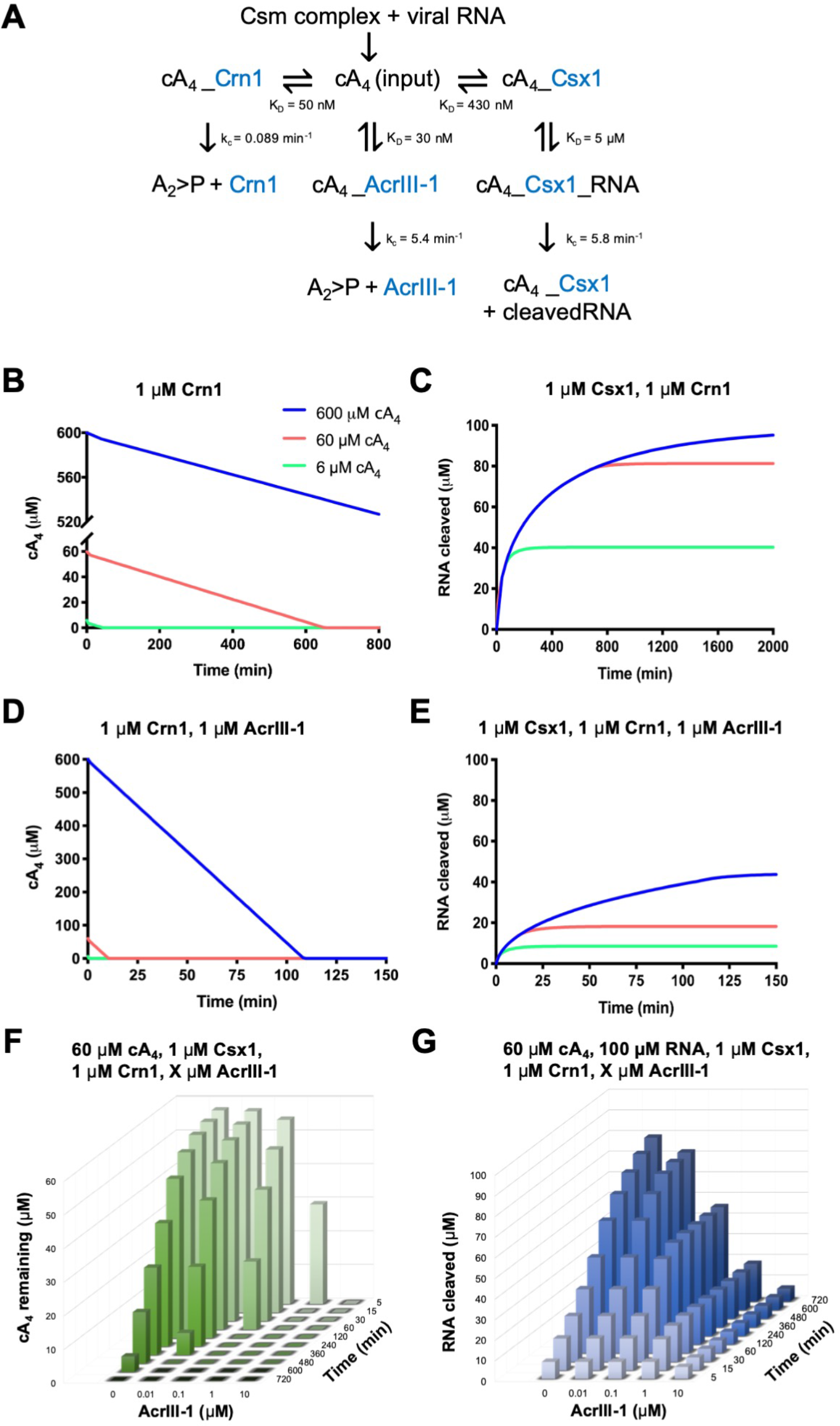
Modelling of *S. solfataricus* antiviral signalling. **A.** Schematic showing kinetic and equilibrium parameters inserted into the KinTek Global Kinetic Explorer software for modelling the type III CRISPR defence illustrated in figure 1. Underscores connecting two variables indicate their relationship in a complex. cA_4_, cyclic tetra-adenylate; Crn1, CRISPR ring nuclease 1; AcrIII-1, viral ring nuclease anti-CRISPR SIRV1 gp29; Csx1, CRISPR ancillary ribonuclease; A_2_>P, di-adenylate containing 2’,3’ cyclic phosphate (product of cA_4_ cleavage). Progress curves depict (**B**) cA_4_ (600 μM, blue; 60 μM, salmon; 6 μM, green) cleavage by 1 μM Crn1 alone or together with 1 μM AcrIII-1 (**C**). Panels **D** and **E** depict RNA (100 μM at start) cleavage by Csx1 (1 μM) in the presence of cA_4_ concentrations as colour coded previously, and its attenuation by 1 μM Crn1 or 1 μM AcrIII-1, respectively. **F** is a 3D plot visualising the concentration of cA_4_ remaining (from 60 μM at start) in the presence of 1 μM Crn1 and varying amounts of AcrIII-1 across a range of doubling endpoints. **G** is a 3D plot visualising concentration of RNA (100 μM at start) cleaved by Csx1 in the presence of 60 μM cA_4_, 1 μM Crn1 and varying amounts of AcrIII-1.

**Table 1.** Summary of experimentally derived parameters used for modelling the type III antiviral signaling pathway. Enzyme concentrations were set initially at 1 μM, based on published studies of transcript levels (Ortmann *et al.*, 2008, Wurtzel *et al.*, 2010), but were varied during modelling to assess the influence of enzyme concentration on RNA cleavage.

| | assumed<br>cellular<br>conc (μM) | K<sub>D</sub> cA<sub>4</sub><br>(nM) | K<sub>D</sub> RNA<br>(μM) | k<sub>c</sub> RNA<br>(min<sup>-1</sup>) | k<sub>c</sub> cA<sub>4</sub><br>(min<sup>-1</sup>) |
| --- | --- | --- | --- | --- | --- |
| **Csx1** | 1 | 430 | 5 | 5.8 | - |
| **Crn1** | 1 | 50 | - | - | 0.089 |
| **AcrIII-1** | 1 | 30 | - | - | 5.4 |

Strikingly, when AcrIII-1 was introduced to the model, even 600 μM cA_4_ was degraded rapidly and Csx1 activity was strongly supressed (Figure 6D, E). This is consistent with our previous biochemical comparison of Crn1 and AcrIII-1 (Athukoralage *et al.*, 2020), and confirms the qualitative difference between the cellular and viral ring nucleases, leading to fundamentally different outcomes on infection. The concentration of AcrIII-1 within *S. solfataricus* cells during infection is not known. In order to determine its correlation to Csx1 deactivation we varied AcrIII-1 concentration in the model and simulated RNA cleavage (Figure 6F, G). We ascertained the AcrIII-1 levels required to significantly decrease cleavage of 100 μM RNA by Csx1, by first challenging 60 μM cA_4_ with increasing AcrIII-1 concentrations. AcrIII-1 concentrations as low as 100 nM slowed RNA cleavage dramatically, allowing no more than 30% of the RNA to be degraded. In contrast, when challenged with 600 μM cA_4_, ≥1 μM AcrIII-1 was required to notably impact Csx1 deactivation (Figure 6-figure supplement 2). This illustrates that the level of AcrIII-1 required to attenuate antiviral signaling is governed by the concentration of cA_4_ generated during the immune response. Therefore, during infection, a positive correlation between AcrIII-1 concentration and viral transcript levels would be required for continued escape from type III CRISPR defence – a reasonable assumption.

Further, by varying the concentration of enzymes involved in the antiviral signalling pathway, we examined the effects of increasing Csx1 and the subsequent burden on Crn1 and AcrIII-1 to downregulate its activity. In particular, we found that 1 μM AcrIII-1 alongside 10 μM Crn1 was inadequate to degrade 60 μM cA_4_ and deactivate 10 μM Csx1 in a manner mirroring speedy abrogation of RNA cleavage when equimolar concentrations of the three enzymes were present. The requirement for greater concentrations of ring nucleases, despite the unaltered rate of RNA cleavage upon increasing Csx1 concentration, is likely a reflection of the competition between Csx1 and ring nucleases for cA_4_ governed by the relevant equilibrium binding constants. Hence increased Csx1 expression may be used to counter AcrIII-1 inhibition of Csx1 activity and could additionally be employed to drive cells to dormancy or death, if the Crn1 concentration was held significantly below that of Csx1.

## DISCUSSION

### Signal amplification in type III CRISPR defence

In this study, we used biochemical data to build a kinetic model of the type III CRISPR antiviral signalling pathway within *S. solfataricus* cells and examined the capacity of CRISPR and anti-CRISPR ring nucleases for its regulation. Quantification of cA_4_ generated by the SsoCsm complex revealed that ~1000 molecules of cA_4_ are made per RNA target, amounting to a concentration of 6 μM in the cell. This large degree of signal amplification ensures that detection of 1 RNA target can generate sufficient amounts of cA_4_ to fully activate the ribonuclease effector protein Csx1, which has a dissociation constant for cA_4_ of 0.4 μM. Given the large signal amplification observed here, it seems likely that some means of cOA degradation, either via self-limiting ribonucleases (Athukoralage *et al.*, 2019, Jia *et al.*, 2019) or dedicated ring nucleases (Athukoralage *et al.*, 2018), will be essential for type III CRISPR systems to provide immunity rather than elicit abortive infection. Indeed, growth arrest has been observed for cOA activated Csm6 during bacteriophage infection (Rostol & Marraffini, 2019). This life or death decision in response to genotoxic stress has also been observed in *S. islandicus*, which becomes dormant upon viral infection and subsequently dies if virus remains in culture (Bautista *et al.*, 2015). In recent years, diverse CRISPR systems have been implicated in abortive infection or cell dormancy. The Type I-F CRISPR system of *Pectobacterium atrosepticum* was found to provide population protection by aborting infection when infected by virulent phage (Watson *et al.*, 2019). Likewise, the in-trans collateral RNA cleavage of *Listeria seeligeri* Cas13a resulted in cell dormancy, providing herd immunity to the bacterial population (Meeske *et al.*, 2019). Similarly, in ecological contexts, it is possible that different multiplicities of viral infection illicit different outcomes from the type III CRISPR response that benefit either the individual cell or the population.

### Cellular and viral ring nucleases reset the system in fundamentally different ways

Biochemical comparison of Crn1 and AcrIII-1 revealed that both enzymes bind cA_4_ with dissociation constants around 40 nM, around 10-fold tighter than observed for Csx1. However, Crn1 is a much slower enzyme. Kinetic modelling of the antiviral signalling pathway confirms that Crn1 is effective only at low levels of viral gene expression, where it has the potential to neutralise the toxicity associated with cA_4_ activated ribonucleases to offer a route for cell recovery without abrogating immunity. In contrast, the much faster reaction kinetics of the anti-CRISPR ring nuclease means it can rapidly deactivate Csx1 and immunosuppress cells even under very high RNA target (and thus cA_4_) levels.

Our modelling suggests that the rapid turnover of cA_4_ by AcrIII-1 over a wide concentration range greatly limits RNA cleavage by deactivating defence enzymes. Therefore, the deployment of AcrIII-1 upon viral infection may not only promote viral propagation but also safeguard cellular integrity until viral release by lysis. Recent studies have uncovered that sequentially infecting phage evade CRISPR defences by exploiting the immunosuppression achieved by Acr enzymes from failed infections (Borges *et al.*, 2018, Landsberger *et al.*, 2018). Further, these immunosuppressed cells have been shown to be susceptible to Acr-negative phage infections, highlighting the complex ecological consequences of supressing CRISPR immunity (Chevallereau *et al.*, 2019). In *Sulfolobus* Turreted Icosahedral virus (STIV), the AcrIII-1 gene *B116* is expressed early in the viral life cycle (Ortmann *et al.*, 2008). Therefore AcrIII-1 accumulation in the cell, possibly from early expression by unsuccessful viruses may, as our models demonstrate, favour the success of latter viral infections. Type III CRISPR systems also conditionally tolerate prophage (Goldberg *et al.*, 2014), and unsurprisingly, AcrIII-1 is found in a number of prophages and integrative and conjugative elements. In these cases, constitutively expressed AcrIII-1 may further immunocompromise cells, and sensitise them to infection by phage otherwise eradicated by type III CRISPR defence. In the ongoing virus-host conflict, while increasing Csx1 concentration may allow better immunity when faced with AcrIII-1, upregulating AcrIII-1 expression in cells will undoubtedly offer viruses an avenue for counter offence.

It should be noted that the type III CRISPR locus of *S. solfataricus* contains a number of CARF domain proteins and their contribution to immunity has not yet been studied. In particular, the CARF-family putative transcription factor Csa3 appears to be involved in transcriptional regulation of CRISPR loci, including the adaptation and type I-A effector genes, when activated by cA_4_ (Liu *et al.*, 2015, Liu *et al.*, 2017). These observations suggest that the cOA signal may transcend type III CRISPR defence in some cell types by activating multiple defence systems. However, by degrading the second messenger, AcrIII-1 has the potential to neutralise all of these.

### Cyclic nucleotides in prokaryotic defence systems

Cyclic nucleotide-based defence systems are emerging as powerful cellular sentinels against parasitic elements in prokaryotes. Mirroring the role of cyclic GMP-AMP synthase (cGAS) in eukaryotic defence against viruses as part of the cGAS-STING pathway, bacterial cGAS enzymes have recently been discovered that abort infection by activating phospholipases through cGAMP signaling (Cohen *et al.*, 2019). Termed the cyclic-oligonucleotide-based antiphage signaling system (CBASS), a large number of additional cOA sensing effector proteins associated with CBASS loci remain uncharacterised, highlighting great diversity in the cellular arsenal used for defence (Burroughs *et al.*, 2015, Cohen *et al.*, 2019). Furthermore, diverse cyclic dinucleotide cyclases have been identified that generate a range of cyclic nucleotides including cUMP-AMP, c-di-UMP and cAAG, which are also likely to function in novel antiviral signal transduction pathways (Whiteley *et al.*, 2019). Type III systems also generate cyclic tri-adenylate (cA_3_) and cyclic penta-adenylate (cA_5_) molecules. Whereas no signalling role has yet been ascribed to cA_5_, cA_3_ has been demonstrated to activate a family of DNases termed NucC which abort infection by degrading the host genome prior to completion of the phage replication cycle (Lau *et al.*, 2020).

The balance between immunity, abortive infection and successful pathogen replication is likely to be governed by enzymes that synthesise and degrade these cyclic nucleotide second messengers. Just as prokaryotes with type III CRISPR require a means to degrade cOA in appropriate circumstances, eukaryotic cells have enzymes that degrade cGAMP to regulate cGAS-STING mediated immunity (Li *et al.*, 2014). Likewise, while prokaryotic viruses utilise AcrIII-1 to rapidly degrade cA_4_, eukaryotic poxviruses utilise Poxins to subvert host immunity by destroying cGAMP (Eaglesham *et al.*, 2019), and pathogenic Group B *Streptococci* degrade host c-di-AMP using the CndP enzyme to circumvent innate immunity (Andrade *et al.*, 2016). The rate of discovery of new defence pathways and cyclic nucleotide signals is breath-taking. Analysis of the dynamic interplay between enzymes that leads to fluctuations in the levels of these second messengers is therefore of crucial importance if we are to achieve an understanding of these processes.

## METHODS

### Cyclic oligoadenylate (cOA) synthesis and visualisation

Cyclic tetra-adenylate (cA_4_) made per RNA target (0.01, 0.1, 1, 10, 25 or 50 nM) was investigated in a 20 μl reaction volume incubating A26 RNA target or A26 phosphorothioate RNA target (Table 2) with 13.5 μg *Sulfolobus solfataricus* (Sso)Csm complex (~470 nM carrying A26 CRISPR RNA) in Csx1 buffer containing 20 mM MES pH 5.5, 100 mM K-glutamate, 1 mM DTT and 3 units SUPERase•In™ Inhibitor supplemented with 1 mM ATP, 5 nM α-^32^P-ATP and 2 mM MgCl_2_ at 70 °C for 2 h. All samples were deproteinised by phenol-chloroform extraction (Ambion) followed by chloroform (Sigma-Aldrich) extraction prior to separating the cOA products by thin-layer chromatography (TLC). TLC was carried out as previously described (Rouillon *et al.*, 2019). In brief, 1 μl of radiolabelled cOA product was spotted 1 cm from the bottom of a 20 × 20 cm silica gel TLC plate (Supelco Sigma-Aldrich). The TLC plate was placed in a sealed glass chamber pre-warmed at 37 °C containing 0.5 cm of a running buffer composed of 30% H_2_O, 70% ethanol and 0.2 M ammonium bicarbonate, pH 9.2. After TLC the plate was air dried and sample migration visualised by phosphor imaging. For analysis, densiometric signals corresponding to cA_4_ was quantified as previously described (Rouillon *et al.*, 2019).

**Table 2.**
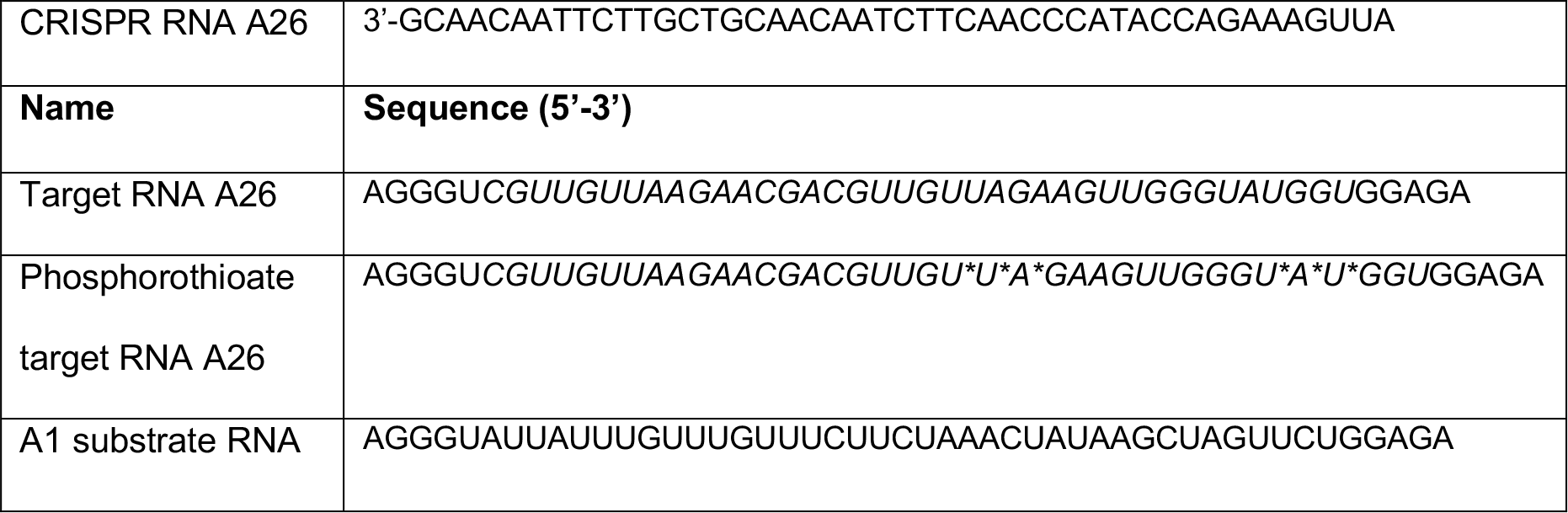
Oligonucleotides. CRISPR RNA A26 is shown 3’ to 5’. Phosphorothioate linkages are indicated with an asterisk. Regions complementary to CRISPR RNA A26 are italicized.

### Generation of α-^32^P-ATP standard curves

cA_4_ synthesis was visualised by incorporation of 5 nM α-^32^P-ATP added together with 0.5 mM ATP at the start of the reaction. Therefore, to calculate the concentration of ATP used for cA_4_ synthesis, α-^32^P-ATP standard curves were generated in duplicate, starting with 5 nM α-^32^P-ATP within a 20 μl volume to represent the densiometric signal corresponding to the complete conversion of 0.5 mM ATP into cOA. Serial two-fold dilutions of 5 nM α-^32^P-ATP and 0.5 mM ATP starting from a 20 μl volume were made and 1 μl of each dilution was spotted on a silica plate and phosphorimaged alongside TLC separating cOA made with varying RNA target concentrations. After phosphorimaging, the densiometric signals of the serial dilutions were quantified, averaged and plotted against ATP concentration starting from 0.5 mM and halving with each two-fold dilution. A line of best fit was then drawn. The concentration of ATP used to synthesise cA_4_ was calculated by entering the densiometric signal of the cA_4_ product into to equation of the line of best fit for the α-^32^P-ATP standard curve. The concentration of cA_4_ generated was derived by dividing the concentration of ATP incorporated by four to account for polymerisation of four ATP molecules to generate one molecule of cA_4_. Finally, the molecules of cA_4_ made per RNA was calculated by dividing the cA_4_ concentration generated by the concentration A26 RNA target used for cOA synthesis.

### Calculation determining the concentration of cA_4_ made when one RNA target is detected within a S. solfataricus cell of ≈ 0.8 μm (0.6-1.0 μm) diameter

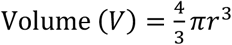 and 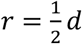

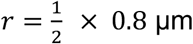

*r* = 0.4 μm

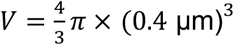

*V* = 0.268 μm^3^

1 μm^3^ = 1 fL

0.268 μm^3^ = 0.268 fL = 2.68 × 10^−13^ mL

1 mole of RNA = 6.022 × 10^23^ molecules of of RNA

1 molecule of RNA = 1 ÷ 6.022 × 10^23^ = 1.661 × 10^−24^ moles of RNA

As ~1000 molecules of cA_4_ is made per 1 molecule of RNA

1.661 × 10^−24^ moles × 1000 = 1.661 × 10^−21^ moles of cA_4_

Concentration (M) = moles / Volume (L)

1.661 × 10^−24^ moles ÷ 2.68 × 10^−16^L = 6.20 × 10^−6^ M or 6. 20 μM cA_4_

### Electrophoretic mobility shift assays to determine cA_4_ equilibrium binding constants

~20 nM radioactively-labelled cA_4_ generated using the SsoCsm was incubated with increasing concentrations of Csx1 (0.01, 0.05, 0.10, 0.20, 0.40, 0.60, 0.80, 1,0, 2.0, 4.0, 8.0, 10.0, 20.0 μM protein dimer) in buffer containing 20 mM Tris-HCl pH 7.5, 150 mM NaCl, 2 mM MgCl_2_ supplemented with 2 μM Ultrapure Bovine Serum Albumin (Invitrogen) for 10 min at 25 °C. A reaction volume equivalent of 20 % (v/v) glycerol was then added prior to loading the samples on a 15 % polyacrylamide, 1 × TBE gel. Electrophoresis was carried out at 28 °C and 250 V. Gels were phosphor imaged overnight at −80 °C. For investigating RNA binding, 50 nM 5’-end radiolabelled and gel purified A1 RNA was incubated with Csx1 variant H345N (0.01, 0.10, 1.0, 5.0, 10.0, 20.0 μM protein dimer) in the presence or absence of 20 μM cA_4_ for 15 min at 40 °C. To examine cA_4_ binding by Crn1, ~10 nM radiolabelled SsoCsm cA_4_ was incubated with Sso2081 (0.01, 0.05, 0.10, 0.20, 0.40, 0.60, 0.80, 1,0, 2.0, 4.0, 8.0, 10.0, 20.0 μM protein dimer) on ice for 15 min before gel electrophoresis as described above but at 300V and at 4 °C. cA_4_ binding by AcrIII-1 was examined by incubating ~10 nM radiolabelled SsoCsm cA_4_ with SIRV1 gp49 H47A (0.001, 0.01, 0.02, 0.03, 0.04, 0.05, 0.06, 0.08, 0.10, 1.0, 10.0, 20.0 μM protein dimer) for 10 min at 25 °C before gel electrophoresis at 30 °C as described above. For analysis densiometric signal corresponding to cA_4_ bound protein was quantified. The densiometric count corresponding to cA_4_ bound to 20 μM Csx1 dimer was used to represent 100% binding and densiometric counts from other lanes were normalised to this value within each replicate. Error of the 100% bound (20 μM Csx1 dimer) densiometric count was derived by calculating the area adjusted count for each replicate and then the standard deviation of their mean, reporting the standard deviation as a fraction of the mean set as 100% bound.

### Single turnover kinetics of RNA cleavage by Csx1

Single turnover kinetic experiments were carried out by incubating Csx1 (2 μM dimer) with A1 RNA (50 nM) in the presence of cA_4_ (20 μM) in Csx1 buffer at 50 °C. This temperature was set somewhat below the normal growth temperature of *Sulfolobus* (75 °C) to allow rate calculations, consistent with previous studies (Athukoralage *et al.*, 2018, Athukoralage *et al.*, 2020). Control reactions with no protein and with protein and RNA in the absence of cA_4_ were included. 10 ul reaction aliquots were quenched by adding to phenol-chloroform and vortexing at 15 s intervals up to 2 min and at 3, 4, 5, 6, and 8 min. Deproteinised products were run on a 7 M urea, 20 % acrylamide, 1 X TBE gel at 45 °C as previously described (Rouillon *et al.*, 2019), and phosphorimaged overnight at −80 °C. Experiments were carried out in triplicate. For analysis the fraction of substrate RNA cut compared to the RNA only control was plotted and fitted to an exponential rise equation as previously described (Rouillon *et al.*, 2019).

### Modelling antiviral signalling and its control by ring nucleases

Modelling was carried out using the KinTek Explorer™ 8 software package (Johnson, 2009), which is available from (https://kintekcorp.com/software). Experiments were modelled and simulated using kinetic and equilibrium paramters detemined experimentally as described in Figure 5A. The following steps were inserted to generate the model:

cA4 + Csx1 = cA4_Csx1
cA4_Csx1 + RNA = cA4_Csx1_RNA
cA4_Csx1_RNA = cA4_Csx1_cleavedRNA (irreversible)
cA4_Csx1_cleavedRNA = cA4_Csx1 + cleavedRNA
cA4 + Crn1 = cA4_Crn1
cA4_Crn1 = A2 + Crn1 (irreversible)
cA4 + Vrn = cA4_Vrn
cA4_Vrn = A2 + Vrn (irreversible)

Simulations were carried out varying cA_4_ concentration (6, 60 and 600 μM) while Csx1, Crn1 (Sso2081) and AcrIII-1 concentration was fixed at 1 μM dimer, or varied depending on the simulation, with total substrate RNA in the cell fixed at 100 μM.

---

**Figure 2-figure supplement 1:**
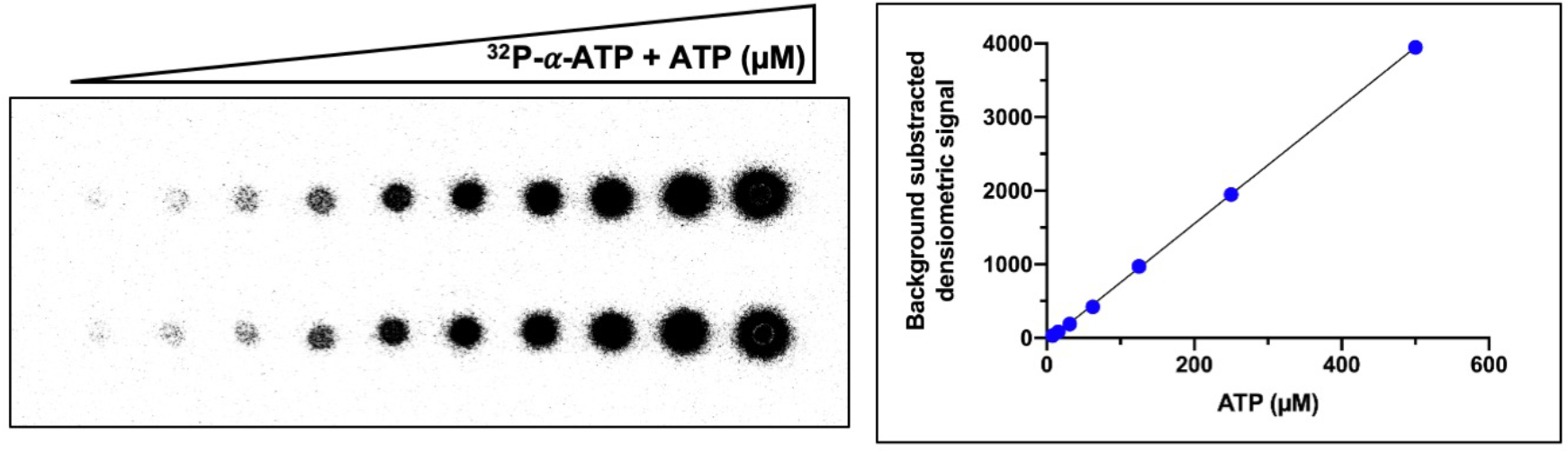
Example of ATP standard curve used to determine the concentration of ATP converted to cyclic tetra-adenylate (cA_4_). Left-hand side panel shows duplicate serial dilution of 32P-α-ATP (5 nM) and ATP (500 μM) mix spotted (1 μl) on a thin-layer chromatography (TLC) plate. The right-hand side panel is a plot of the densiometric signal quantified from the TLC plate after phosphorimaging. The mean densiometric signal is plotted and errors bars showing the standard deviation are plotted but not visible due to their scale. The densiometric signal corresponding to cA_4_ was compared to the standard curve to determine the concentration of ATP converted. Duplicate standard curves were carried out for each replicate assay examining cA_4_ synthesis.

**Figure 6-figure supplement 1:**
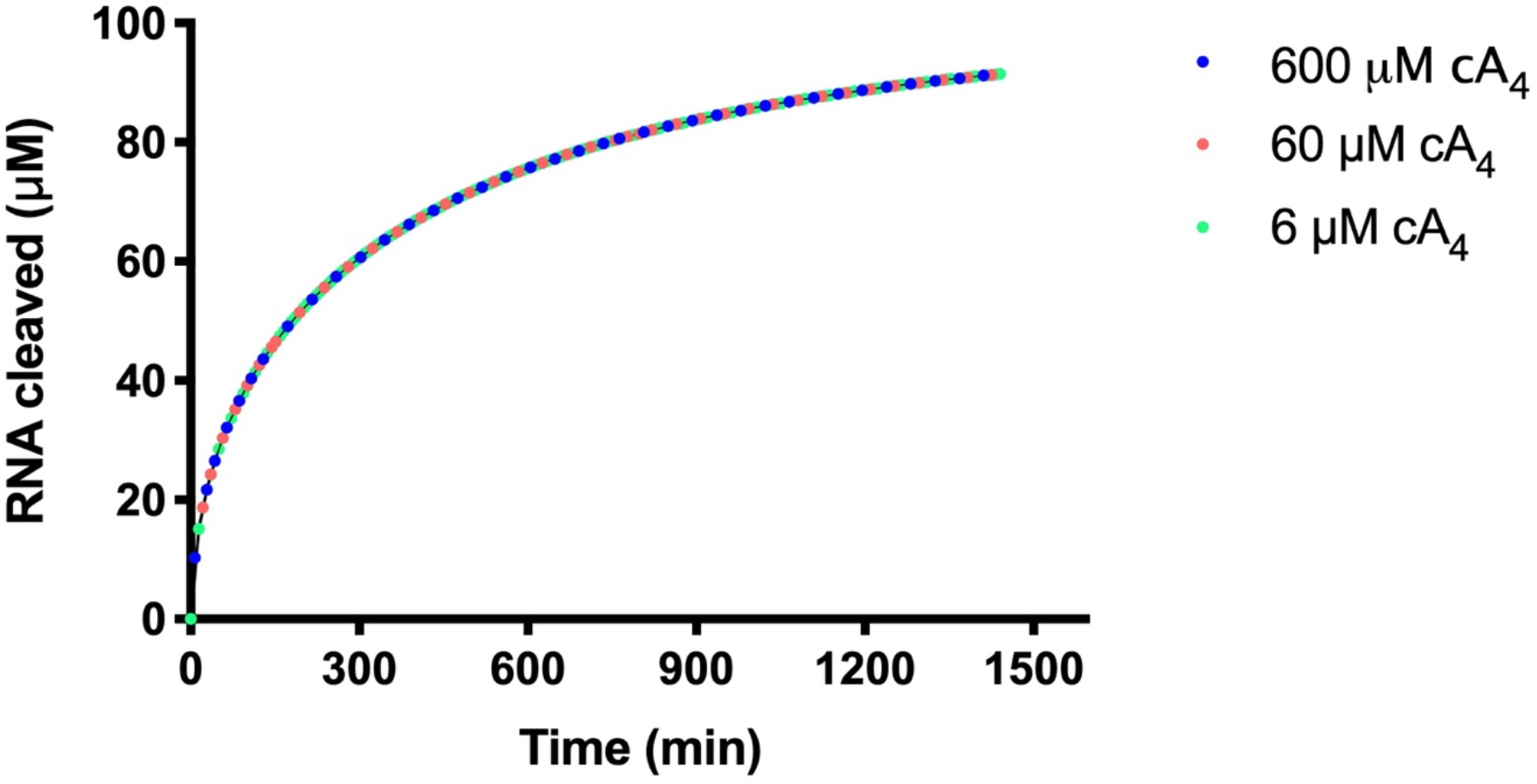
RNA cleavage by Csx1 in the presence of varying cA_4_ concentrations. Plot of RNA (100 μM) cleaved by 1 μM Csx1 in the presence of 6, 60 or 600 μM cA_4_ and no ring nuclease. Identical amounts of RNA are cleaved when cA_4_ is in excess of Csx1 concentration and all the RNA present is eventually degraded.

**Figure 6-figure supplement 2:**
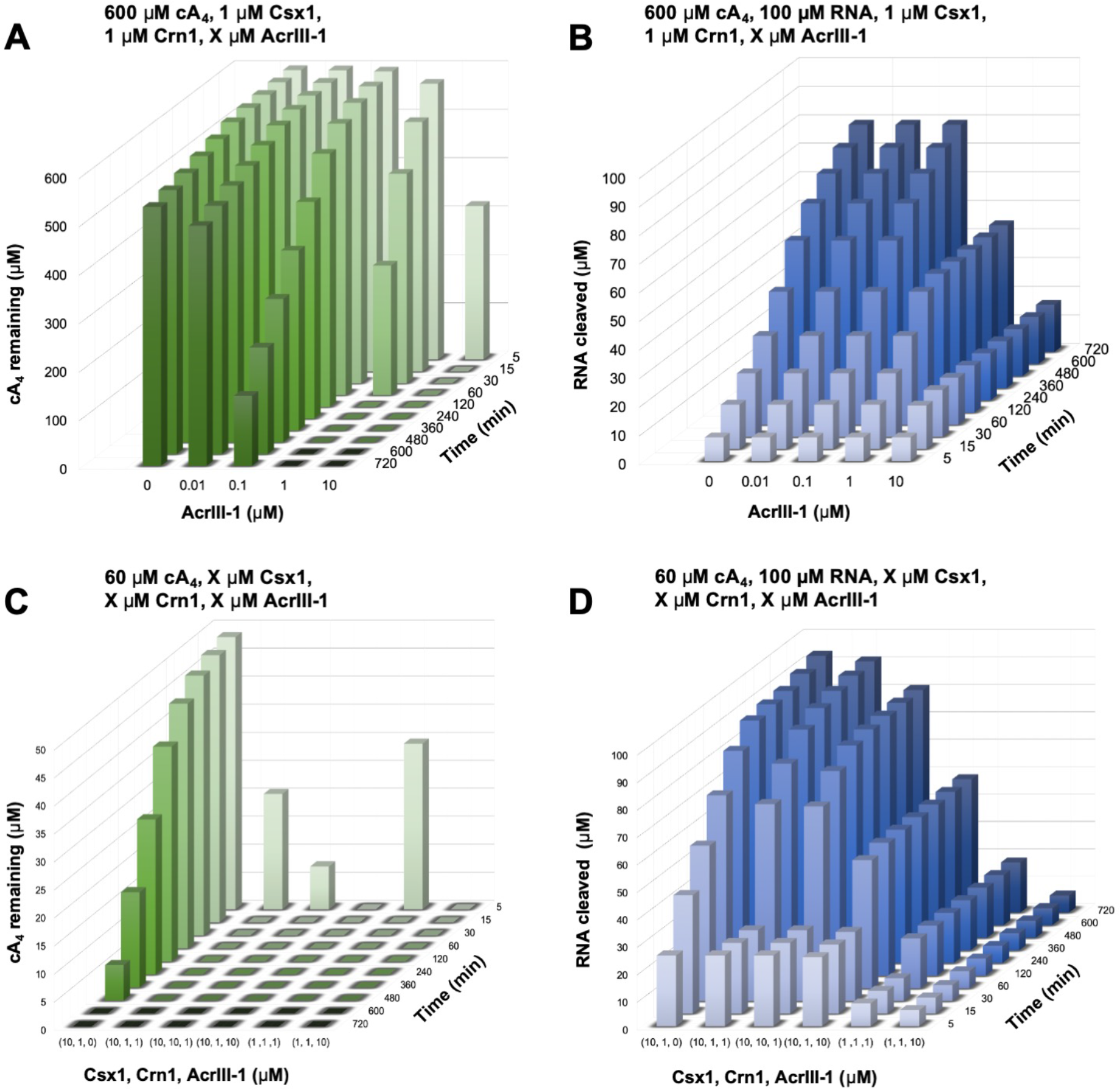
RNA and cA_4_ degradation under varied cA_4_ and enzyme concentrations. **A.** 3D plot of cA_4_ remaining when 600 μM cA_4_ is challenged with varying concentrations of AcrIII-1 in the presence of Csx1 (1 μM) and Crn1 (1 μM). **B.** Effect of varying AcrIII-1 on RNA (100 μM) cleavage over time, under conditions as in A. **C.** Effect of varying Crn1 and/or AcrIII-1 on cleavage of 60 μM cA_4._ **D.** RNA (100 μM) cleaved when Csx1 concentration is varied together with Crn2 and/or AcrIII-1, under conditions as in C.

## Acknowledgments

This work was supported by a grant from the Biotechnology and Biological Sciences Research Council (Grant REF BB/S000313/1 to MFW) and the Wellcome Trust (Grant 210486/Z/18/Z to CMC)

## Author contributions

Januka S. Athukoralage, Investigation, Methodology, Formal analysis, Visualisation, Writing-original draft preparation, Writing-review and editing; Shirley Graham, Methodology and editing; Christophe Rouillon, Methodology and editing; Sabine Grüschow, Methodology and editing; Clarissa M. Czekster, Methodology, Visualisation and editing; Malcolm F. White, Conceptualisation, Formal analysis, Supervision, Project administration, Funding acquisition, Writing-original draft preparation, Writing-review and editing.

